# The evolution of latitudinal ranges in reef-associated fishes: heritability, limits, and inverse Rapoport’s rule

**DOI:** 10.1101/2020.11.02.365700

**Authors:** Marcio R. Pie, Raquel Divieso, Fernanda S. Caron, Alexandre C. Siqueira, Diego R. Barneche, Osmar J. Luiz

## Abstract

**Aim:** Variation in the size and position of geographical ranges is a key variable that underlies most biogeographical patterns. However, relatively little is known in terms of general principles driving their evolution, particularly in the marine realm. In this study we explore several fundamental properties regarding the evolution of reef fish latitudinal ranges, namely the degree of similarity in range size between ancestor and descendant lineages (i.e. phylogenetic signal); the evolution of range limits; and the latitudinal distribution of range sizes, particularly with respect to Rapoport’s rule.

**Location:** Global.

**Taxon:** Reef-associated fishes.

**Methods:** We integrate data on the latitudinal distribution and evolutionary history of 5,071 reef fish species with phylogenetic comparative methods to assess the level of phylogenetic signal in latitudinal range size, low- and high-latitude limits, and range midpoints, and to estimate rates of evolution of those traits. Finally, we test whether latitudinal ranges become smaller near the equator, as predicted by Rapoport’s rule, using phylogenetic generalized least squares.

**Results:** There were varying levels of phylogenetic signal in latitudinal range size, low- and high-latitude limits, and range midpoints. Despite these differences, latitudinal midpoints were consistently shown to have the highest phylogenetic signal among all measured geographic features. Interestingly, the position of high-latitude limits in general evolved at substantially faster rates than their low-latitude counterparts. Finally, we confirm for the first time the existence of an inverse Rapoport’s rule in reef-associated fishes using phylogenetic comparative methods. Indeed, mean latitudinal range size of tropical species is nearly twice the size of their temperate counterparts (2067±1431 km vs. 1168±725 km, respectively).

**Main conclusions:** We uncovered several congruent patterns in phylogenetic signal and rates of evolution of latitudinal ranges, despite vastly disparate biogeographical distributions and ecological differences between the studied fish lineages. Such broad congruence across different taxa and oceans, as well as with previous data from terrestrial environments, suggests that the observed patterns might represent general principles governing geographical range evolution.

## INTRODUCTION

Some of the most pervasive patterns in the distribution of life on earth, such as the latitudinal gradient in species diversity (Willig, Kaufman & Stevens, 2003; Hillebrand, 2004; Mittelbach et al., 2007) and the differences in faunal composition between regions (Ficetola, Mazel & Thuiller, 2017; Cowman et al., 2017), are fundamentally determined by the evolution of the position and limits of geographical ranges of different organisms. For instance, phylogenetic niche conservatism posits that the ancestral niche can determine the regions and habitats that a clade can colonize, and those in which it will persist in the face of environmental change (Wiens & Donoghue, 2004). Therefore, understanding the evolution of geographical ranges has far implications for many areas in ecology, conservation and evolution, from biogeography to the response of species to climate change. In this study we focus on three fundamental properties of geographical distributions: the degree of similarity in latitudinal range size between ancestor and descendant lineages, the evolution of range limits, and the geographical distribution of range sizes using reef-associated fishes as a model system. Reef fishes are an ideal group for studying patterns and processes in range size in the marine realm. They are conspicuous members of shallow-water reef ecosystems, facilitating the surveys that provide presence-absence data needed to quantify range sizes (Kulbicki et al., 2013). Furthermore, reef fishes are among the most diverse groups of vertebrates in the world, exhibiting extraordinary taxonomic breadth and endemism (Cowman et al., 2017). Species within this group also vary greatly in both their range size and in many of the potential biological traits that may influence range size (Ruttenberg & Lester, 2015).

The seminal paper of Jablonski (1987) demonstrated that sister species of Cretaceous mollusks exhibited correlation in their range sizes, thus suggesting that there may be heritability in this emergent trait. However, the generality of species-level “heritability” of emergent traits in general, and of geographical range size in particular, is still unclear. For example, although studies on terrestrial taxa have shown mixed support for range size heritability (e.g. Ricklefs & Latham, 1992; Blackburn & Gaston, 1996; Waldron, 2007), some of these inferences might have been biased by the low statistical power of the early methods used to test for species level heritability (more recently referred to as phylogenetic signal, see Revell, Harmon & Collaret, 2008), such as variance partitioning. On the other hand, even though model-based methods were more successful in detecting phylogenetic signal in range size (Cardillo, 2015; Pie & Meyer, 2017), they also found substantial differences among lineages depending on their biological characteristics (e.g. Freckleton, Harvey & Pagel, 2002, see also Ricklefs & Latham, 1992). Interestingly, although the original study of Jablonski (1987) involved marine mollusks, no study to date tested for range size heritability in extant marine species (but see Roy et al., 2009).

One cannot understand the evolution of geographical range size without first recognizing what determines its limits (Gaston, 2009; Sexton et al., 2009). This issue was well presented by Merriam (1894): “What naturalists wish to know is not how species are dispersed, but how they are checked in their efforts to overrun the earth”. The study of range limits is particularly intriguing in the ocean, given that there are few hard biogeographical barriers that inhibit species dispersal when compared with terrestrial habitats (Palumbi, 1994, but see Gaines et al., 2009). Within marine habitats, the proximate mechanisms driving the distribution of species might involve some combination of environmental tolerances (Rezende & Bozinovic, 2019; Deutsch, Penn & Seibel, 2020; Marshall et al., 2020), biological interactions (Longo et al., 2019), species-level traits that have the potential to impact species’ distributions by influencing colonization ability and/or post-colonization survival or persistence (Luiz et al., 2013), and the impact of ocean currents on larval dispersal (e.g. Gaylord & Gaines, 2000; Mora & Sale, 2002; Álvarez-Noriega et al., 2020). While these proximate mechanisms have been scrutinized in the macroecological literature, the ultimate (evolutionary) mechanisms driving range limit evolution in marine species are still poorly known (but see Sunday et al. 2012). In terrestrial organisms, tolerance to heat is largely conserved across lineages, whereas tolerance to cold varies between and within species, a pattern that was interpreted as evidence for hard physiological boundaries constraining the evolution of tolerances to high temperatures (Araújo et al., 2013, see also Qu & Wiens, 2020). Indirect evidence of environmental tolerances can be obtained by recording the minimum and maximum latitudes of occurrence of a given species, given the widespread association between geographical distributions and physiological processes, particularly in marine fish (Stuart-Smith et al., 2015; Stuart-Smith, Edgar & Bates, 2017; Waldock et al., 2019; Dahlke et al., 2020). Given that many climatic variables vary consistently with latitude, such limits can serve as a proxy for the corresponding environmental limits. Indeed, Pie & Meyer (2017) found that, for terrestrial vertebrates, rates of evolution of high-latitude limits were 1.6-4 times faster than low-latitude limits, suggesting that different mechanisms drive the evolution of “warm” and “cold” range boundaries. The extent to which this pattern is shared with marine organisms is currently unknown.

Rapoport’s rule states that species that are characteristic of high-latitude locations have greater latitudinal range than species which inhabit low-latitude locations (Rapoport, 1982). Therefore, it could potentially provide a mechanism to explain the latitudinal gradient in species richness (Stevens, 1989, but see Šizling et al., 2009). Although evidence for Rapoport’s rule has been obtained for several terrestrial taxa, its generality has been called into question (Gaston, Blackburn & Spicer, 1998) because the prediction of Stevens (1989) was based on an incomplete model of species distributions and did not take into account the extent and geometry of environmental gradients. Species distribution models that simulate range dynamic of species along latitudinal gradients can generate traditional Rapoport-like, inverse, or flat gradients in range size, depending on the actual gradients in mean and seasonality of climatic and other factors (Tomašových et al. 2015). Moreover, it has rarely been assessed in marine organisms. For instance, the few studies to date testing Rapoport’s rule for marine fishes found that, not only the rule did not apply, but also that ranges actually became larger towards the equator (Rohde, Heap & Heap, 1993; Macpherson & Duarte, 1994; Macpherson, Hastings & Robertson, 2009, but see Fortes & Absalão, 2010). However, it is important to note that, given the often significant phylogenetic signal in geographical ranges (e.g. Cardillo, 2015; Pie & Meyer, 2017), it is possible that the obtained results might have been affected by phylogenetic nonindependence between species, potentially biasing both the direction and the significance of the estimated relationships. To the best of our knowledge, no study to date tested for the existence of Rapoport’s rule in marine fishes using phylogenetic comparative methods.

In this study we provide a comprehensive assessment of the evolution of geographical ranges in reef fishes. In particular, we focus on three main questions: (1) is there significant phylogenetic signal in their latitudinal range sizes? (2) do low- and high-latitude range limits evolve at different rates? and [3) do patterns of reef fish latitudinal range size variability support Rapoport’s rule?

## MATERIALS AND METHODS

### Data collection

Latitudinal limits were obtained from the presence-absence dataset of Rabosky et al. (2018). This dataset consists of global occurrences of marine species in geographic grid cells of 150 km^2^, and it was built through the AquaMaps algorithm (Ready et al., 2010). From this dataset, we filtered those species belonging to families defined by Siqueira *et al* (2020) as associated with reef habitats *sensu lato* (not only coral reefs). To overcome the challenges of defining reef fishes as an operational unit (Bellwood & Wainwright, 2002), Siqueira *et al* (2020) used a systematic selection criterion. They first assessed the full list of families that had been previously characterized as having reef-associated species (Bellwood & Wainwright, 2002). Subsequently, a list of all valid species within those families was created by using the ‘rfishbase’ (Boettiger *et al* 2012) R package. Finally, data from ‘rfishbase’ was used to calculate the proportion of reef-associated species in each family, and only families with more than 20% of reef-associated species were kept in the dataset. We used this dataset that included all valid species within 65 selected fish families (see Siqueira *etal* 2020). It is important to highlight here that, by including all valid species within the selected families, we kept some species in our dataset that were not necessarily reef-associated nor limited to tropical latitudes (proportion of reef-associated species provided in Supplementary table S1). This was important to avoid biases in calculating phylogenetic signal and rates of range evolution. Our final dataset contained geographical range data for 5,071 fish species.

Our inferences rely fundamentally on the validity of the obtained distribution data. Ideally, one would rather rely on occurrence data from sampling sites throughout the areas in which the studied species could potentially be found, yet that is rarely the case for most biogeographical datasets. However, several important strategies were used by Rabosky et al. (2018, from which we obtained the distribution data) to ensure that the results would reflect real patterns in species distributions. First, they transformed the modelled distribution data on each cell based on a fixed threshold of 0.5, such that the probability of occurrence of each species in a given cell was at least 50%. Second, they used expert opinions, when available, to refine the projected distributions by truncating the predicted distributions in light of museum data, known biogeographical barriers and specialized literature on particular data. Therefore, although the biogeographical database has limitations, it is likely to reflect the underlying variation in distribution patterns of the studied species. We recognize that the geographical range—the projection across geographical space of the locations where a given species can be found—is a complex phenomenon with temporal and spatial heterogeneity in its manifestation. However, we focused on three particular aspects of their range, namely the highest, lowest and midpoint latitudinal coordinates, as a first approximation, given that these properties of the geographical ranges are most likely to reflect interspecific variation in climatic conditions and physiological tolerances. Whenever a distribution spanned the equator, the lowest and highest limits were determined based on their absolute values, such that the limit closer to the equator was the lowest limit and vice-versa, as in Pie and Meyer (2017).

To perform our comparative analyses, we used the phylogenetic trees produced by Siqueira et al. (2020). This set of 100 phylogenies was produced by the taxonomic imputation of missing species into the backbone tree of Rabosky et al. (2018) using the TACT algorithm (Chang, Rabosky & Alfaro, 2020). In this approach, birth-death-sampling estimators are used across ultrametric phylogenies to estimate branching times for unsampled taxa, with taxonomic information to compatibly place new taxa onto a backbone phylogeny.

### Analyses

*We* estimated the phylogenetic signal of the latitudinal midpoint, as well as the high- and low-latitude limits of reef fishes using Pagel’s λ (Pagel, 1999) and Blomberg et al.’s *K* (2003) using the *phylosig* function in ‘phytools’ 0.7-47 (Revell, 2012). More specifically, we tested whether a model with an estimated λ provides a better fit than a simpler alternative where λ=0, whereas *K* estimates were based on 1,000 resamplings of the original data. Given that these approaches have slightly different assumptions, we used both to ensure that the estimates of phylogenetic signal are consistent. Latitudinal range size data were Intransformed prior to the analyses. We used phylogenetic generalized least squares (PGLS, Freckleton et al., 2002) to test the relationship between the latitudinal range (difference between high and low latitude limits as the response variable) and the absolute latitudinal position (predictor variable) using the *pgls* function in ‘caper’ 1.0.1 (Orme et al., 2008). We estimated the rate of evolution of each geographic feature based on the σ^2^ parameter under a Brownian Motion (BM) model using the *fitContínuous* function in ‘geiger’ 2.0.7 (Harmon et al., 2008). Differences in rates of evolution of low- and high-latitude limits were tested using the *mvBM* function in ‘mvMORPH’ 1.1.3 (Clavel, Escarguel & Merceron, 2015). This test contrasts the fit of a model in which low- and high-latitude limits share the same rates against an alternative where each limit has a separate rate using likelihood ratio tests. To account for phylogenetic uncertainty, we repeated all analyses on each of the 100 alternative topologies and report the median of the estimates of each parameter and model likelihood. However, given that the phylogenetic imputation used to provide those topologies might have introduced noise in the inferred parameters, we repeated all analyses using a smaller phylogeny (2,585 tips) that included only those species for which genetic data were available (derived from Rabosky et al. 2018). We refer to these datasets as *expanded* and *reduced,* respectively. The random imputation of species based on taxonomy can also bias the inferred evolutionary signal of distribution features. Therefore, the tests of phylogenetic signal were performed using the reduced dataset only. In each of the above-mentioned analyses, we pruned the trees to estimate separately the evolutionary rates and phylogenetic signal for different families and orders, as long as the resulting subtrees had at least 20 tips. Finally, untransformed latitude values were used to compute lower and upper limits because otherwise it would be difficult to compare species present in both hemispheres with those present in only one. However, sensitivity analyses excluding transequatorial species provided qualitatively similar results and will not be explored further here (see Supplementary Material, Tables S10, S11, S12 and S13). All analyses were carried out in R 4.0.2 (R Core Team, 2020).

## RESULTS

The distribution of latitudinal ranges in the studied fish species follows the typical pattern found in terrestrial organisms, with an approximately lognormal distribution with a left skew (Figure 1), as opposed to a strong right skew in untransformed data (not shown). The truncation on the right-hand side of the distribution is probably the result of a geometric constraint in terms of the size of ocean basins, given that the largest possible range (i.e. the natural logarithm of the difference between the maximum and minimum latitudes across all studied species) would be ≈ 4.7. Despite this hard boundary condition, the average latitudinal ranges of the studied lineages tended to be relatively broad, with many species approaching the largest latitudinal range classes. There were varying levels of phylogenetic signal in latitudinal range size, low- and high-latitude limits and range midpoints, both at the order (Figure 2) and at the family levels (Figure S1). Despite these differences, range midpoints consistently showed the highest levels of phylogenetic signal, both for Pagel’s λ and Blomberg et al.’s *K*(Tables S2, S3).

**Figure 1.**
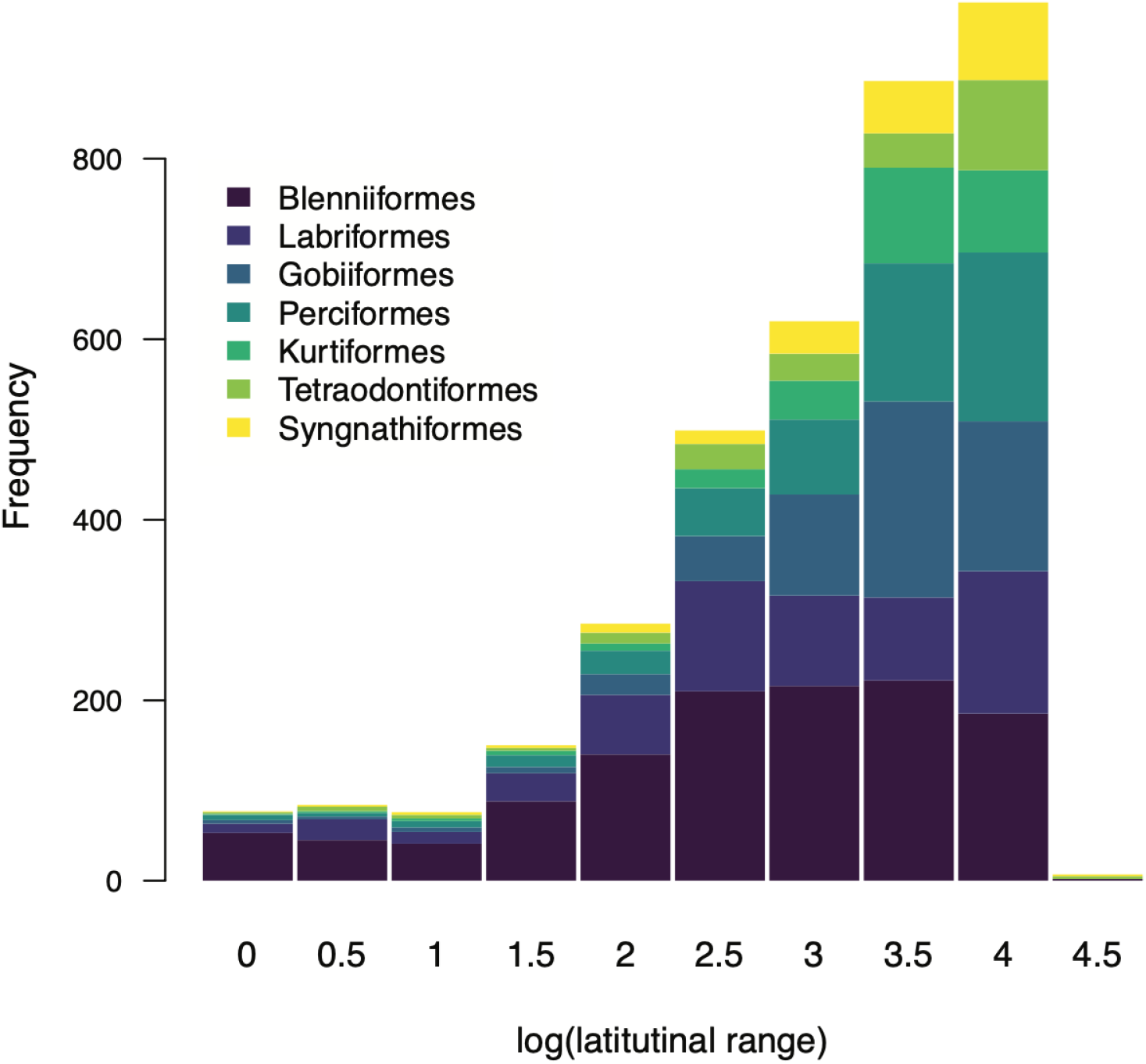
Distribution of log-transformed latitudinal ranges of the seven most species-rich marine fish orders in our dataset. The largest possible range (i.e. the natural logarithm of the difference between the maximum and minimum latitudes across all studied species) would be ≈ 4.7.

**Figure 2.**
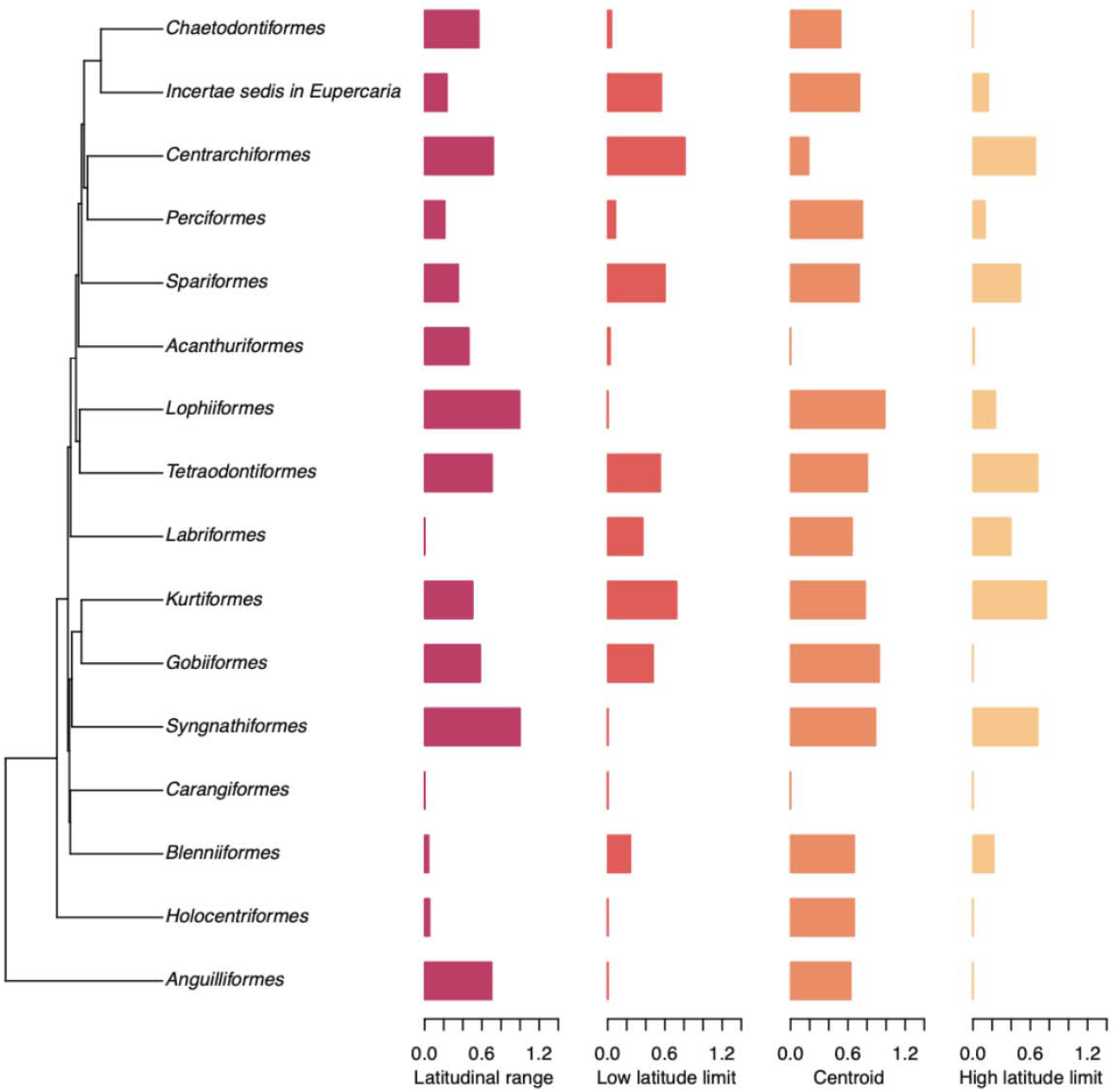
Phylogenetic signal, according to Pagel’s λ (Pagel 1999), of properties of geographical ranges of reef-associated fish orders. More information on how this tree was inferred, as well as corresponding node support values, can be obtained in Rabosky et al. (2018) and Siqueira et al. (2020). Details on estimates, corresponding likelihoods and p-values are indicated in Table S2.

Results of statistical tests for variation in rates of evolution between high- and low-latitude limits for different fish orders are shown in Table 1. Significant rate differences were detected in 17 and 13 out of the 19 and 16 tested orders for the expanded and reduced datasets, respectively. These significant differences predominantly involved faster rates in the location of high-latitude limits (16 out of 17 and 10 out of 12 for the expanded and reduced datasets, respectively). At the family level, significant rate differences were detected in 36 and 17 families out of the 41 and 27 tested for the expanded and reduced datasets, respectively (Table S4). Again, significant differences predominantly involved faster rates in the location of high-latitude limits (33 versus 3 and 13 versus 4 for the expanded and reduced datasets, respectively). Therefore, although not universal, a faster rate of evolution of high-latitude range limits is widespread among reef fish lineages. Finally, contrary to the expectation based on Rapoport’s rule, in general there was a negative relationship between latitude and latitudinal range (Figure 3), which was significant in 11 out of 15 orders (Table 2) and 19 out of 23 families for the reduced dataset (Table S5). There was no case of a positive relationship between latitude and latitudinal range predicted by Rapoport’s rule for any of the tested orders and families. Indeed, when we compared the mean latitudinal range size of exclusively tropical species vs. exclusively temperate species, the former were nearly twice larger (2067±1431 km vs. H68±725 km, respectively).

**Table 1.**
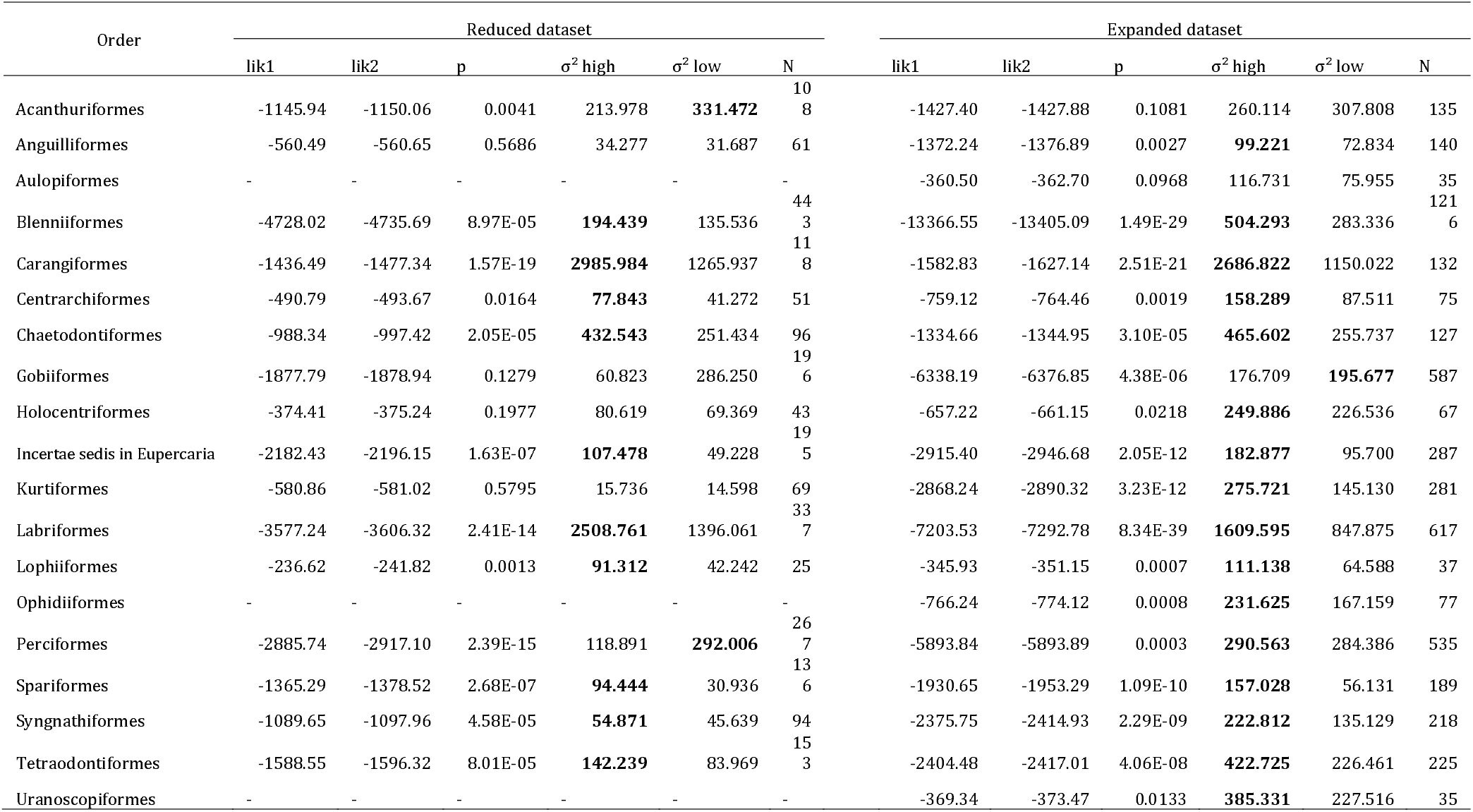
Relative fit of models in which high- and low-latitude range limits of marine reef fish orders evolve at the same or different rates: lik1: log likelihood of multiple-rate model; lik2: likelihood of single-rate model; σ^2^ high and σ^2^ low indicate estimated rates of evolution for high- and low-latitude range limits, respectively; N: number of analyzed species. Values for the expanded dataset indicate medians across 100 alternative topologies. Bold-face rates are significantly different.

**Figure 3.**
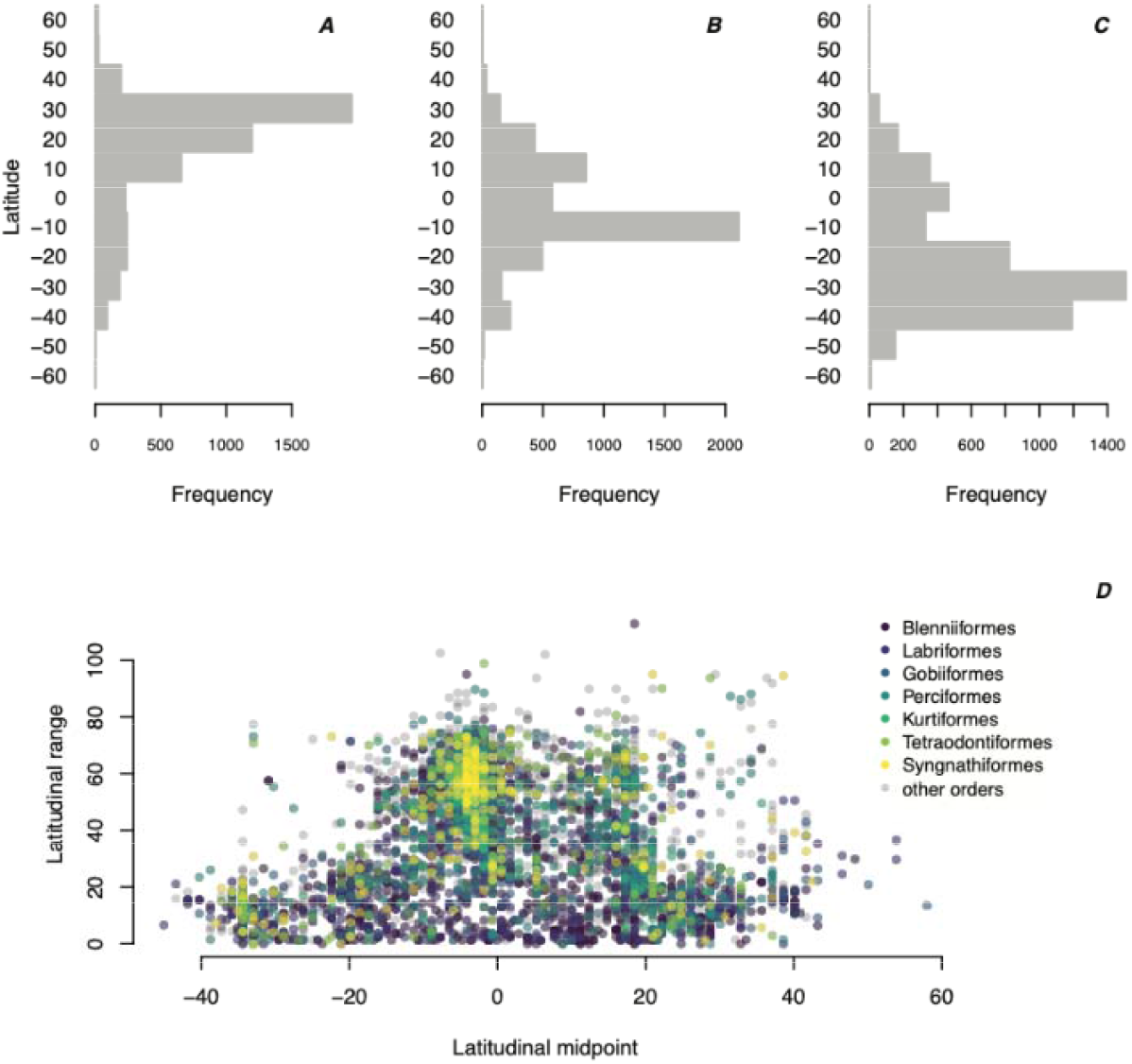
Latitudinal variation in the northern limit (A), midpoint (B), and southern limit (C) of marine reef fishes, as well as the latitudinal distribution of their corresponding latitudinal range sizes.

**Table 2.**
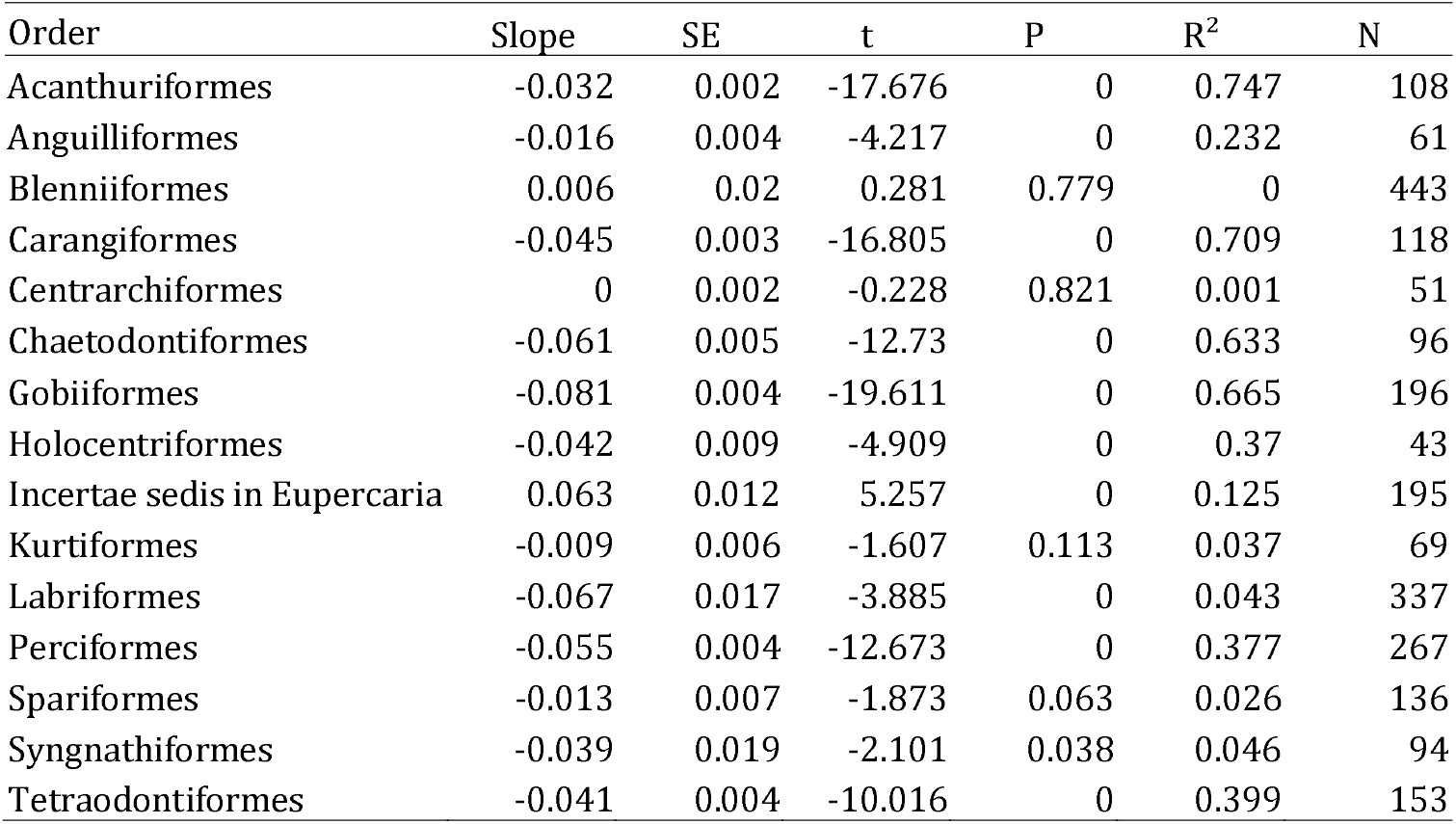
Phylogenetic generalized least squared analyses of the relationship between absolute latitude and latitudinal range of different reef fish orders.

| Order | Slope | SE | t | P | R<sup>2</sup> | N |
| --- | --- | --- | --- | --- | --- | --- |
| Acanthuriformes | -0.032 | 0.002 | -17.676 | 0 | 0.747 | 108 |
| Anguilliformes | -0.016 | 0.004 | -4.217 | 0 | 0.232 | 61 |
| Blenniiformes | 0.006 | 0.02 | 0.281 | 0.779 | 0 | 443 |
| Carangiformes | -0.045 | 0.003 | -16.805 | 0 | 0.709 | 118 |
| Centrarchiformes | 0 | 0.002 | -0.228 | 0.821 | 0.001 | 51 |
| Chaetodontiformes | -0.061 | 0.005 | -12.73 | 0 | 0.633 | 96 |
| Gobiiformes | -0.081 | 0.004 | -19.611 | 0 | 0.665 | 196 |
| Holocentriformes | -0.042 | 0.009 | -4.909 | 0 | 0.37 | 43 |
| Incertae sedis in Eupercaria | 0.063 | 0.012 | 5.257 | 0 | 0.125 | 195 |
| Kurtiiformes | -0.009 | 0.006 | -1.607 | 0.113 | 0.037 | 69 |
| Labriformes | -0.067 | 0.017 | -3.885 | 0 | 0.043 | 337 |
| Perciformes | -0.055 | 0.004 | -12.673 | 0 | 0.377 | 267 |
| Spariformes | -0.013 | 0.007 | -1.873 | 0.063 | 0.026 | 136 |
| Syngnathiformes | -0.039 | 0.019 | -2.101 | 0.038 | 0.046 | 94 |
| Tetraodontiformes | -0.041 | 0.004 | -10.016 | 0 | 0.399 | 153 |

It is important to note that, given that the Atlantic and Indo-Pacific ocean basins were influenced by different histories and environmental gradients, we repeated our analyses separately for the evolutionary rate tests, and considered whether species were present in either (or both) ocean basins as a covariate in PGLS analyses. Rate tests provided qualitatively similar results (Table S6 for orders, Table S7 for families). Likewise, the PGLS for different families (Table S8) and orders (Table S9) did not show a significant effect of ocean basin on the relationship between latitude and latitudinal range except for the order Perciformes and the family Scorpaenidae. However, when the corresponding data were visualized (Figure S2), it becomes clear that the significant effect of ocean basin on Perciformes is mostly due to species with very large range sizes that occur in both basins whose range sizes are not related to latitude (Figure S2). Although there was also a significant effect of ocean basin on the scorpaenid range data, it was mostly reflected in different slopes of relationships that were still negative, thus corroborating that none of the studied fish taxa showed evidence for a positive relationship between range size and latitude that is expected based on Rapoport’s rule.

## DISCUSSION

In this study we provide the most comprehensive assessment of the evolution of latitudinal ranges of marine fishes to date. In particular, we uncover several general principles governing geographical range evolution in reef-associated fishes, namely (1) there was stronger phylogenetic signal in latitudinal midpoint than in latitudinal limits; (2) high-latitude range limits evolve substantially faster than low-latitude range limits; and (3) latitudinal range sizes tend to become larger in species whose midpoints are closer to the equator.

The detection of phylogenetic signal in traits related to geographical ranges might provide insight into important mechanisms driving their evolution. For instance, if closely related species show range overlap, they could experience similar conditions that would lead to parallel expansions and contractions in their geographical ranges when faced with climate change, as seems to be the case in birds (Mouillot & Gaston, 2009). However, our results indicate that not all properties of reef fish latitudinal ranges share similar levels of phylogenetic signal. Rather, range midpoints tended to show higher phylogenetic signal than either latitudinal range or limits. These differences suggest that the evolution of latitudinal ranges takes place at a relatively constant rate, whereas range size and limits might experience stronger heterotachy, thus obscuring patterns of phylogenetic autocorrelation. These results are particularly intriguing when compared to the estimates of evolutionary rates (Table 1, Table S4). The combination of relatively lower phylogenetic signal in high-latitude range limits and their higher evolutionary rates is not obvious, given that higher rates do not necessarily entail lower phylogenetic signal (Revell et al., 2008). Rather, these results provide a scenario in which tropical reef fish gradually change the midpoint position and size of their latitudinal ranges, whereas range limits (particularly those at higher latitudes) show a faster and more heterogeneous rate of evolution. One potential mechanism to explain this pattern is the observation that higher latitudes have experienced more variation in temperatures through time (Zachos et al. 2001; Siqueira et al., 2016; Gaboriau et al., 2019). An alternative scenario posits that environmental stresses drive high-latitude limits, whereas biotic interactions would be more important at low latitudes, although evidence in support for this scenario is still scarce (Sax 2008). More detailed studies, particularly using more precise geographical distribution data, as well as more detailed phylogenetic comparative methods of range evolution, might provide important insight into the extent to which this scenario is representative of reef fish range evolution in general.

We confirmed previous studies suggesting an inverse Rapoport rule for marine fishes (Rohde et al., 1993; Macpherson & Duarte, 1994). In particular, the latitudinal range of tropical species was nearly twice that of their temperate counterparts, on average. It is important to note that Macpherson (2003) and Fortes & Absalão (2010) actually found patterns consistent with Rapoport’s rule in smaller and larger datasets than ours, respectively, of marine fishes. However, they used the method proposed by Stevens (1989), which compares the mean latitudinal range of all species within each 5° latitudinal band within the latitudinal gradient. As indicated by Rohde et al. (1993), in this method the same species are counted multiple times as they are distributed across different latitudinal bands, leading to statistical issues of nonindependence (see also Rohde & Heap, 1996; Ribas & Schoereder, 2006). Although the midpoint method used in our study might lead to some level of bias towards the inverse Rapoport’s rule (Connolly, 2009), this effect is more likely in the case of a large fraction of species with very broad ranges, which is not the case in our dataset. For comparison, we carried out additional analyses comparing range sizes of species that were exclusively tropical or temperate (thus avoiding the problem of nonindependence of the Stevens method), and our conclusions remained unaltered. Interestingly, an inverse Rapoport’s rule was also found for marine prosobranch gastropods living on the shelves of the western Atlantic and eastern Pacific Oceans (Roy et al., 1998) and for marine bivalves in the western and eastern Pacific and western Atlantic coasts (Tomašových et al., 2015).

The reverse Rapoport’s rule appears to be common among marine organisms because of the nonlinearity of the latitudinal gradient of temperature in the sea, with much weaker spatial variation in annual minimum, mean and maximum daily temperature at low than at mid latitudes. These effects may be a general factor shaping latitudinal gradients in range size, because at the global scale, as shallow thermal gradients within the tropics and steep thermal gradients at higher latitudes also characterize some terrestrial environments (Tomašových et al., 2015). There are three potential mechanisms to explain the inverse Rapoport’s rule in marine species. First, the less steep environmental gradients at low latitudes could allow for broader geographic ranges in the tropics, whereas the steeper temperature gradients at high latitudes can prevent the complete occupation of those regions (Pintor et al., 2015; Tomašových et al., 2015), as has been suggested in the case of marine bivalves (Tomašových et al., 2015). Bivalve thermal range size does not peak in the tropics because tropical species achieve broad latitudinal ranges even when they are thermally specialized (Tomašových & Jablonski 2016), so that the climatic variability hypothesis is sufficient to explain latitudinal distribution of thermal range size but is insufficient to explain distribution of geographical range sizes. Moreover, available evidence suggests that the relationship between the thermal tolerance breadth of a species and the latitude at which it is typically encountered is very weak for marine ectotherms (Sunday et al. 2011, see also Parmesan et al. 2005). An alternative mechanism is the mid-domain effect (MDE), which would be expected even in the absence of climatic gradients (Colwell & Hurt, 1994). In particular, the MDE predicts that the random distribution of species, while following the geometrical constraints imposed by the outer limits of the latitudinal domain, would follow a quasi-parabolic gradient in species diversity with most species being found in the center of the domain (Colwell & Hurt, 1994, see also Šizling et al. 2009). Under the MDE, a species with a range midpoint at high latitudes would necessarily have a smaller range (otherwise its latitudinal midpoint would be closer to the equator). On the other hand, lower latitudes could include both large- and small-range species, as they would not be limited geometrically, as indeed was shown by our results (Figure 3D). It is important to note that, based on our own results, the MDE alone cannot explain the similarity in closely-related species in their latitudinal positions, given that the detected phylogenetic signal in range properties means that a simple lineage shuffling process is not realistic. Finally, secondary biodiversity hotspots driven by parapatric speciation processes, as well as overlaps between tropical and subtropical species, could also be contributing to this pattern (Pinheiro et al., 2018).

To what extent do latitudinal ranges in marine and terrestrial organisms evolve according to the same rules? These differences have been the focus of intense research in recent years (e.g. Webb 2012, Pinsky et al. 2019, Schumm et al. 2019). However, other than the fact that latitudinal ranges of marine organisms are commonly considered much larger than those of terrestrial organisms (Gaston, 2003), there have been few attempts to assess the extent to which geographical range evolution might differ between these two environments. We argue that, despite the obvious differences between marine and terrestrial life, their geographical ranges actually share several important characteristics, namely (1) approximately lognormal distributions with a left skew (Gaston, 2003; Macpherson, 2003; Mora & Robertson, 2005; Figure 1); (2) phylogenetic signal in geographical range properties (Cardillo, 2015; Pie & Meyer, 2017; Figure 2, Figure S1, Tables S2, S3); (3) asymmetry in evolutionary rates between high- and low-latitude limits (Pie & Meyer, 2017; Table 1, Table S4). The congruence of these patterns across taxa and realms indicates that they might indeed represent general principles regarding geographical range evolution that transcend differences between land and ocean habitats. However, there were some intriguing taxa that seem to depart from this expectation, which indicate that these principles are general, but not universal. Therefore, one of the main contributions of our study is that we can now actually recognize those taxa as exceptional, and we can now seek to understand why they are not constrained by the same mechanisms as other taxa.

There are some important caveats with respect to our analyses. For instance, we have not explicitly considered differences among species with respect to depth, which might be an important confounding factor for marine fishes (e.g. Macpherson & Duarte, 1994). However, in the case of reef fishes, depth does not seem to play an important role in explaining interspecific variation in range size (Luiz et al., 2012, 2013). It is also important to note that our evolutionary rate estimates depend fundamentally on (1) the accuracy of the phylogenetic relationships between the studied species and (2) the extent to which distribution ranges evolve according to a Brownian motion model. Formally assessing phylogenetic accuracy is a challenging task, particularly for large-scale phylogenies with thousands of species, in which the level of topological error might vary across lineages based on the level of phylogenetic and taxonomic sampling. Such inaccuracies have been shown to affect downstream analyses (e.g. Title & Rabosky, 2019). We concede that some level of error is present in the phylogenetic relationships used in the present study, particularly when using phylogenetic imputation (Chang et al., 2020). However, we believe that such inaccuracies are unlikely to alter our conclusions for three main reasons. First, any error in the analyzed phylogenies would lead to undirected random noise as opposed to differentially overestimating high-latitude limits in relation to low-latitude limits. Indeed, such error would actually cause the estimated rate differences to be conservative. Second, the consistency in the results across fish taxa of widely different ecologies and biogeographical distributions (and with terrestrial organisms as well) would be unlikely due to chance alone. Finally, the sensitivity analyses carried out in our study, varying the phylogenies in both the expanded and reduced datasets, indicates that our conclusions are largely unaffected by the topological variation in the underlying trees. With respect to the adequacy of Brownian motion (BM) as a model of range evolution, we can most certainly say that it is not. For instance, BM assumes that species are identical in their trait values at the time of speciation, which would be unrealistic for all but some specific cases of sympatric speciation. Also, BM is unbounded, whereas ranges are bounded both by continents and by the poles. Nevertheless, our goal was not to provide a perfect model of range size but rather a first approximation to estimate rates of range size evolution. At present, there are no adaptations of BM-like processes to model specifically the evolution of geographical ranges. We hope that our results, particularly the asymmetry in the evolutionary rates between low- and high-latitude limits, will become an important component of such models in the future.

## ACKNOWLEDGMENTS

This paper was developed in the context of the National Institutes for Science and Technology (INCT) in Ecology, Evolution and Biodiversity Conservation, supported by MCTIC/CNPq (proc. 465610/2014-5) and FAPEG (proc. 201810267000023). MRP was partially funded by a grant from CNPq/MCT (301636/2016-8). We thank B. Hugueny, H. Pinheiro, and an anonymous reviewer for valuable comments on the manuscript. No permits were required for the present study.

## Supplementary material

**Figure S1.** Phylogenetic signal, according to Pagel’s λ (Pagel 1999), of properties of geographical ranges of reef fish families. Details on estimates, corresponding likelihoods and p-values are indicated in Table S2.

**Figure S2.** Relationship between latitudinal range size (ln-transformed) and latitudinal position for the clades for which Ocean basin had a significant effect, based on PGLS analyses. We only tested taxa that had at least 10 tips in each of the groups.

**Table S1.** Families of marine fish used in the study and their respective proportion of species associated with reefs.

**Table S2.** Tests of phylogenetic signal of latitudinal range size, lower latitude, midpoint and higher latitude limits for different reef fish families.

**Table S3.** Tests of phylogenetic signal of latitudinal range size, lower latitude, midpoint and higher latitude limits for different reef fish orders.

**Table S4.** Relative fit of models in which high- and low-latitude range limits of marine reef fish families evolve at the same or different rates: lik1: log likelihood of multiple-rate model; lik2: likelihood of single-rate model; σ^2^ high and σ^2^ low indicate estimated rates of evolution for high- and low-latitude range limits, respectively; N: number of analyzed species. Values for the expanded dataset indicate medians across 100 alternative topologies. Bold-face rates are significantly different.

**Table S5.** Phylogenetic generalized least squares analyses of the relationship between absolute latitude of the range midpoint and latitudinal range of different reef fish orders

**Table S6.** Relative fit of models in which high- and low-latitude range limits of marine reef fish orders evolve at the same or different rates, analyzing separately each ocean basin: lik1: log likelihood of multiple-rate model; lik2: likelihood of single-rate model; σ^2^ high and σ^2^ low indicate estimated rates of evolution for high- and low-latitude range limits, respectively; N: number of analyzed species. Values for the expanded dataset indicate medians across 100 alternative topologies. Bold-face rates are significantly different.

**Table S7.** Relative fit of models in which high- and low-latitude range limits of marine reef fish families evolve at the same or different rates, analyzing separately each ocean basin: lik1: log likelihood of multiple-rate model; lik2: likelihood of single-rate model; σ^2^ high and σ^2^ low indicate estimated rates of evolution for high- and low-latitude range limits, respectively; N: number of analyzed species. Values for the expanded dataset indicate medians across 100 alternative topologies. Bold-face rates are significantly different.

**Table S8.** Phylogenetic generalized least squares analyses of the relationship between absolute latitude of the range midpoint and latitudinal range of different reef fish orders using ocean basin as a covariate.

**Table S9.** Phylogenetic generalized least squares analyses of the relationship between absolute latitude of the range midpoint and latitudinal range of different reef fish families using ocean basin as a covariate.

**Table S10.** Relative fit of models in which high- and low-latitude range limits of marine fish orders [only for non-transequatorial species) evolve at the same or different rates: lik1: log likelihood of multiple-rate model; lik2: likelihood of single-rate model; σ^2^ high and σ^2^ low indicate estimated rates of evolution for high- and low-latitude range limits, respectively; N: number of analyzed species. Values for the expanded dataset indicate medians across 100 alternative topologies. Bold-face rates are significantly different.

**Table S11.** Relative fit of models in which high- and low-latitude range limits of marine fish families [only for non-transequatorial species) evolve at the same or different rates: lik1: log likelihood of multiple-rate model; lik2: likelihood of single-rate model; σ^2^ high and σ^2^ low indicate estimated rates of evolution for high- and low-latitude range limits, respectively; N: number of analyzed species. Values for the expanded dataset indicate medians across 100 alternative topologies. Bold-face rates are significantly different.

**Table S12.** Relative fit of models in which high- and low-latitude range limits of marine fish orders [only for non-transequatorial species) evolve at the same or different rates, analyzing separately each ocean basin: lik1: log likelihood of multiple-rate model; lik2: likelihood of single-rate model; σ^2^ high and σ^2^ low indicate estimated rates of evolution for high- and low-latitude range limits, respectively; N: number of analyzed species. Values for the expanded dataset indicate medians across 100 alternative topologies. Bold-face rates are significantly different.

**Table S13.** Relative fit of models in which high- and low-latitude range limits of marine fish families [only for non-transequatorial species) evolve at the same or different rates, analyzing separately each ocean basin: lik1: log likelihood of multiple-rate model; lik2: likelihood of single-rate model; σ^2^ high and σ^2^ low indicate estimated rates of evolution for high- and low-latitude range limits, respectively; N: number of analyzed species. Values for the expanded dataset indicate medians across 100 alternative topologies. Bold-face rates are significantly different.

## BIOSKETCH

**Marcio R. Pie** is an evolutionary biologist at the Universidade Federal do Paraná, Curitiba, Paraná, Brazil. He is interested macroecology, macroevolution, phylogenetic comparative methods, and molecular ecology. He is interested in uncovering general principles regarding the evolution of geographical ranges and their relationship with climatic niches. Author contributions: M.R.P., R.D., F.S.C., A.C.S., D.B. and O.J.L. conceived the ideas; A.C.S., D.B. and O.J.L. compiled the data; M.R.P., R.D. and F.S.C. analyzed the data; and M.R.P. led the writing.

